# Species richness, habitat association and odonata diversity of the south-western region of Bangladesh

**DOI:** 10.1101/252890

**Authors:** Sajjad Hossain Tuhin, Md Kawsar Khan

## Abstract

Odonata survey was conducted throughout the south-western region of Bangladesh, concentrated on eight districts and the Sundarban (the largest mangrove forest of the world) from August 2014 to August 2016. A total of 50 species under 30 genera belonging to six families were recorded during the study period. Among the 50 species, 31 species belong to Anisoptera whereas 19 species were recorded from the Zygoptera suborder. Libellulidae and Coenagrionidae were most dominant Anisoptetran and Zygopteran families with 28 and 17 species respectively. One Zygoptera species *Mortonagrion varalli* was newly added to the Odonata fauna of Bangladesh. Among surveyed habitats, species richness was highest in marshland with 38 species followed by 34 species in the lakeside. On the other hand, odonata diversity was lowest in the mangrove area. Our study shows a positive correlation between environmental variables (such as average rainfall, relative humidity, and average temperature) and species richness in the study area. Species richness is highest during monsoon and lowest in winter.

## 1. Introduction

Odonates (dragonflies and damselflies) are one of the earliest winged insects developed in Permian period (Kalkman et al. 2008) and distributed all over the world except Antarctica (Silsby 2001; Grimaldi and Engel 2005; Trueman 2007). Although dragonflies are highly distributed in diverse ecological niches, they are very sensitive to the alteration of their habitats. Hence dragonflies are considered as indicators of freshwater ecosystem (Watson et al. 1982; Brown 1991; Martín and Maynou 2016). Besides ecological indicator, dragonflies and damselflies are extensively studied in evolutionary and ecological research (Córdoba-Aguilar, 2008). At present, 5740 species of Odonata known from diverse ecological niches throughout the world (Subramanian 2009).

Odonates are considered freshwater insects because their females lay egg on water or submerged plants and the larval development occurs under water (Hornung & Rice 2003). Unlike the larva, the adults are aerial. However, their foraging and reproductive success depends heavily on the water reservoirs. Hence odonates assemblage is higher in aquatic habitats (Oppel 2005). Besides water reservoirs, odonata diversity varies in different climatic zones. Similar to other insects order, majority of the dragonfly species inhabits the tropical and subtropical climatic zones (Dumont 1991). Compare to other regions, Indo-Malayan is one of the most diversified region and habitat of the maximum endangered dragonflies (Clausnitzer et al. 2009). Bangladesh being located in the Indo-Burma biodiversity hotspot zone possesses high odonata diversity. Along with the geographical variation, seasonal variation like temperature, humidity and rainfall influences the species richness of dragonflies. Bangladesh has six seasons with warm and wet summer, monsoon and autumn from April to September. Temperature starts to fall after September and the late monsoon, winter and spring are dry and cold although temperature barely goes down below 10°C.

Bangladesh has one of the rich habitat for odonatan diversity because it’s geographical locationand abundance of waterbodies. However, very few studies have been done to annotate the Odonata fauna of Bangladesh. In 2011, Chowdhury and Mohiuddin listed 96 species of Odonates from the eastern region while Khan (2015b) reported 76 species from the northeastern part of Bangladesh. In recent years, a few species have been added to the Bangladeshi Odonata fauna and at present 105 species is known from Bangladesh (Khan 2015a; Khan 2017). However, till date odonata survey has been focused mainly on the eastern region while odonata survey is required in the other parts of Bangladesh especially the southwestern region which has diversified water reservoirs.

Administratively, southwestern part of Bangladesh is mainly under Khulna Division. This division is consists of ten administrative districts and covers a large are of 22,285 Km^2^. The largest tract of mangrove forest of the world, the Sundarbans is also situated under this division and distributed over three districts namely, Khulna, Bhagerhat and Satkhira. Many rivers, canals, ponds and lakes are flowing throughout this parts of Bangladesh. All these freshwater reservoirs are excellent habitats of Odonates. Biswas et al., took first approach to annotate the odonates of this region although their study were limited to Bagerhat district only (Biswas 1980). A long time has been passed since last odonatological survey has been conducted in this region, hence it is important to perform a comprehensive survey to annotate the dragonflies of this region.

In present study, we aim to conduct a broad survey in the south-west region of Bangladesh to document the dragonflies and damselflies of this area. We also intend to update the odonata checklist of this region. We surveyed different water reservoirs to predict the odonata diversity in those water bodies. As seasonal variation influences the odonatan diversity, we also aim to find out how seasonal variation influences the species richness in this region.

## 2. Materials and Methods

### 2.1 Study Site

Khulna division lies between 21°38′ 36″ N to 24°10′ 51″ N and 88°33′ 37″ E to 89°56′ 34″ E (Map 01). The study area is under tropical climatic condition with a mild winter from October to March, hot and humid summer in March to June and humid, warm rainy monsoon in June to October. Temperature varies all the year round; in January and December temperature falls to the lowest at 12-15°C and reaches highest in April-June at 41-45 °C. Daily relative humidity fluctuate between 50-90%, which is lowest in the evening and highest in the morning (Fig. 01). Maximum precipitation is experienced in July with 20-25 days of rain with 368 mm precipitation (Bangladesh Bureau of Statistics 2014).

**Figure 1:**
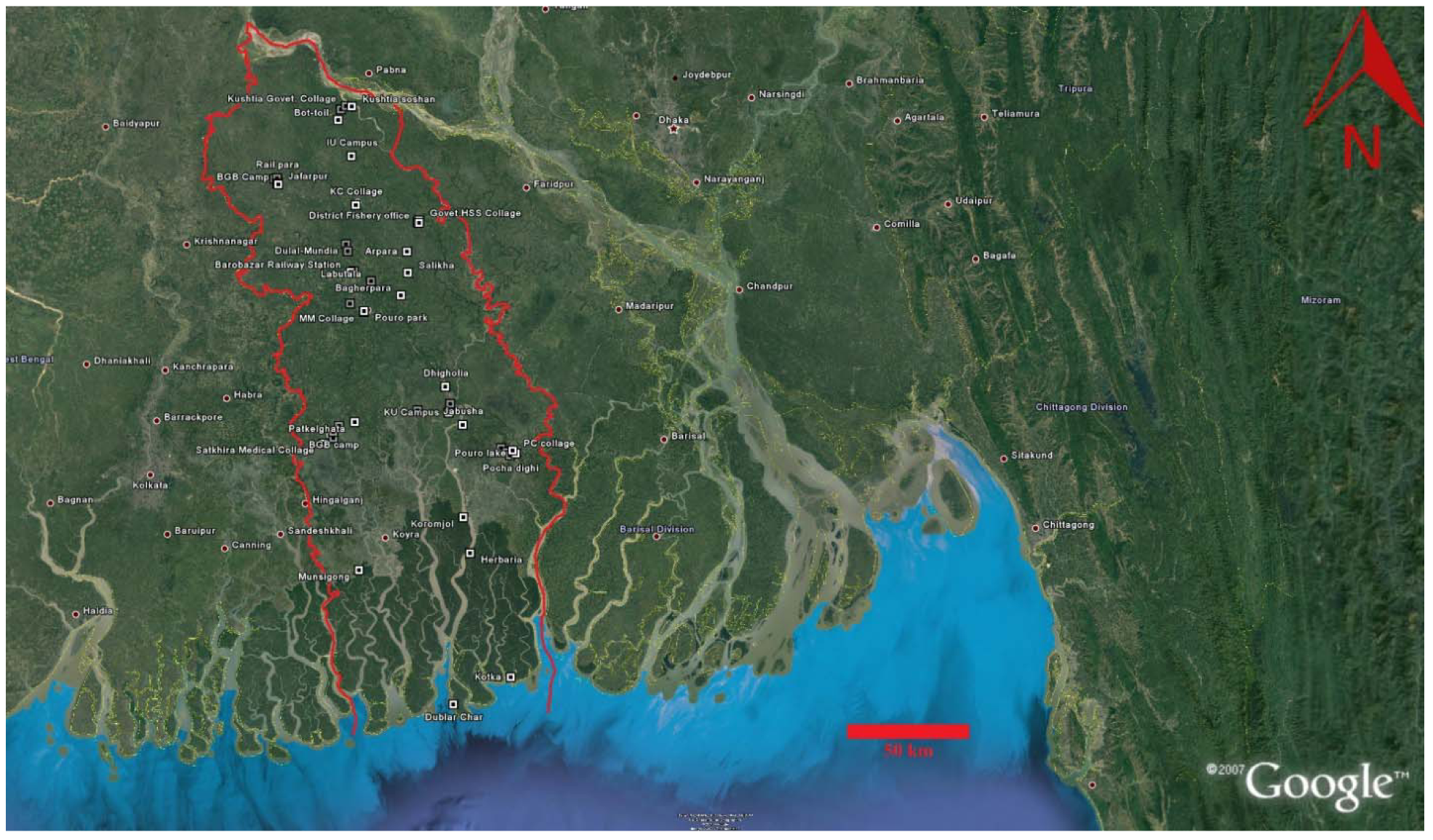
A references map of Southwestern region (Khulna Division) of Bangladesh with survey sites.

### 2.2 Sampling Design

We conducted field works in the south-western region of Bangladesh (concentrated on Khulna Division) from August 2014 to August 2016. We survey eight districts (i.e., Khulna, Kushita, Jessore, Bagherhat, Chuadanga, Satkhira, Magura and Jhinaidha) and the Sundarbans during the study period. We selected those eight districts to cover the whole south-west region of Bangladesh. We selected five sites from every district or zone by considering the accessibility and diversity of the habitat. In total we selected 45 sites in the whole study area (Table 1). We did regular survey in the selected sampling sites under Khulna and Jessore district. We performed weekly survey in Khulna University Campus site in basis, bi-weekly survey in Jabusha; Monthly visit in Boyara and bi-monthly visit in the Pouro Park, M.M Collage, Boart Club and Labutala. We did one or two opportunistic survey in the rest of the sites. We recorded GPS quadrate for all surveyed sites with GPS device (Garmin GPSMAP 76CSx.).

**Table 01.** A list of locations in the south-west region surveyed during the study period.

| Sl. No. | District/Zone | Location | Latitude | Longitude | No of species | Habitat | Date Visited |
| --- | --- | --- | --- | --- | --- | --- | --- |
| 1. | Khulna | KU Campus | 22.80125° N | 89.53466° E | 39 | SU, ML | Weekly visit |
| 2. |  | Boyara | 22.83289°<br>N | 89.54131° E | 18 | FP | Monthly visit |
| 3. |  | Dumuria | 22.80936°<br>N | 89.41360° E | 24 | FP | Bi- Monthly<br>visit |
| 4. |  | Dhigholi<br>a | 22.89416°<br>N | 89.52221° E | 22 | FP | 2 September<br>2016 |
| 5. |  | Jabusha | 22.75812°<br>N | 89.58919° E | 31 | RS, ML | Bi-weekly<br>visit |
| 6. | Jessore | Pouro<br>park | 23.16508°<br>N | 89.20466° E | 16 | LS | Bi- Monthly<br>visit |
| 7. |  | M.M<br>Collage | 23.16234°<br>N | 89.20328° E | 18 | FP | Bi- Monthly<br>visit |
| 8. |  | Bort Club | 23.19148°<br>N | 89.14975° E | 21 | LS | Bi- Monthly<br>visit |
| 9. |  | Labutala | 23.27200°<br>N | 89.23148° E | 29 | FP | Bi- Monthly<br>visit |
| 10. |  | Bagherpa<br>ra | 23.22126°<br>N | 89.34825° E | 14 | ML | 17 June 2016 |
| 11. | Satkhira | Satkhira<br>Govet.<br>Collage | 22.71028°<br>N | 89.08503° E | 19 | FP | 6 September<br>2016 |
| 12. |  | Satkhira<br>Medical | 22.69100°<br>N | 89.04764° E | 14 | ML | 6 September<br>2016 |
|  |  | Collage |  |  |  |  |  |
| 13. |  | BGB<br>camp | 22.73244°<br>N | 89.08453° E | 15 | LS | 6 September<br>2016 |
| 14. |  | Binerpota | 22.75447°<br>N | 89.10717° E | 14 | ML | 6 September<br>2016 |
| 15. |  | Patkelgha<br>ta | 22.76752°<br>N | 89.16900° E | 18 | ML | 6 September<br>2016 |
| 16. | Bagerhat | Satgomuj<br>mosque | 22.67461°<br>N | 89.73788° E | 17 | LS | 9 September<br>2016 |
| 17. |  | Khanjaha<br>n ali<br>majar | 22.66067°<br>N | 89.76032° E | 26 | FP | 9 September<br>2016 |
| 18. |  | PC<br>collage | 22.66543°<br>N | 89.78261° E | 16 | LS | 9 September<br>2016 |
| 19. |  | Pouro<br>lake | 22.65560°<br>N | 89.79535° E | 11 | LS | 9 September<br>2016 |
| 20. |  | Pocha<br>dighi | 22.64870°<br>N | 89.77262° E | 8 | LS | 9 September<br>2016 |
| 21. | Magura | Govet.HS<br>S Collage | 23.48938°<br>N | 89.41914° E | 18 | FP | 15 May 2015 |
| 22. |  | Stadium<br>para | 23.48232°<br>N | 89.41420° E | 12 | SU | 15 May 2015 |
| 23. |  | District | 23.48080° | 89.41904° E | 17 | SU | 15 May 2015 |
|  |  | Fishery office | N |  |  |  |  |
| 24. |  | Arpara | 23.37775°<br>N | 89.37197° E | 11 | RS | 15 May 2015 |
| 25. |  | Salikha | 23.30223°<br>N | 89.37575° E | 16 | ML | 15 May 2015 |
| 26. | Jhinaidha | KC Collage | 23.54539°<br>N | 89.17012° E | 19 | SU | 13 September 2016 |
| 27. |  | Kaligang | 23.40495°<br>N | 89.13213° E | 17 | FP | 13 September 2016 |
| 28. |  | Dulal-Mundia | 23.38004°<br>N | 89.13924° E | 7 | SU | 13 September 2016 |
| 29. |  | Gazi-Kalu Mazar | 23.30836°<br>N | 89.16321° E | 13 | LS | 13 September 2016 |
| 30. |  | Barobazar Railway Station | 23.30488°<br>N | 89.15081° E | 16 | SU | 13 September 2016, 13 August 2017 |
| 31. | Kushita | Kushtia Govet. Collage | 23.90175°<br>N | 89.12837° E | 32 | LS | 3 may 2016, 16 September 2016 |
| 32. |  | Kushtia soshan | 23.89993°<br>N | 89.15211° E | 14 | SU | 16 September 2016 |
| 33. |  | Chourhas | 23.88921°<br>N | 89.10866° E | 9 | SU | 16 September<br>2016 |
| 34. |  | Bot-toil | 23.85114°<br>N | 89.09854° E | 12 | LS | 16 September<br>2016 |
| 35. |  | IU<br>Campus | 23.72105°<br>N | 89.15204° E | 26 | FP | 3 may 2016,<br>16 September<br>2016 |
| 36. | Chuadang<br>a | Chuadan<br>ga Govet.<br>Collage | 23.63979°<br>N | 88.84907° E | 18 | SU | 18 September<br>2016 |
| 37. |  | Rail para | 23.64256°<br>N | 88.85871° E | 23 | SU | 18 September<br>2016 |
| 38. |  | Bus<br>Terminal | 23.63715°<br>N | 88.86063° E | 11 | SU | 18 September<br>2016 |
| 39. |  | Jafarpur | 23.62353°<br>N | 88.85984° E | 14 | ML | 18 September<br>2016 |
| 40. |  | BGB<br>Camp | 23.62049°<br>N | 88.86415° E | 15 | ML | 18 September<br>2016 |
| 41. | Sundarban<br>s | Munsigo<br>ng | 22.23776°<br>N | 89.18557° E | 17 | MF | 3<sup>rd</sup> September<br>2015, 5<br>February<br>2016 |
| 42. |  | Koromjol | 22.42743° | 89.59065° E | 14 | MF | 10 October |
|  |  |  | N |  |  |  | 2014 |
| 43. |  | Herbaria | 22.29961°<br>N | 89.61641° E | 17 | MF | 11 October<br>2014 |
| 44. |  | Dublar<br>Char | 21.75720°<br>N | 89.54883° E | 11 | MF | 14 August<br>2015 |
| 45. |  | Kotka | 21.85464°<br>N | 89.77198° E | 16 | MF | 17 August<br>2015 |

(SU= Semi Urban areas; ML=Marsh Land; V= Vegetation; RS= River Side; LS= Lake Side and MF= Mangrove Forest)

We surveyed the odonates by walking through the different habitat of the study sites (i.e., road sides, canal bank, river bank, pond sides, lake sides, open fields, forest paths, crop fields, grasslands, urban and semi-urban areas) from 8:00 hr to 17:00 hr. We photographed the specimens with various identification keys with a Nikon-3200D camera using Nikkor 55-300mm AF-S DX and Micro-Nikkor 105 mm FX AF lenses. We collected those species which are difficult to identify from visual inspection and photographs by using an insect sweeping net. We collected only few specimens to avoid large distraction of the odonata population. We Identified the Odonates with the help of taxonomic keys provided by Fraser (1933, 1934, 1936), Asahina (1967), Lahiri (1987), Mitra (2002), Subramanian (2005), and Nair (2011) and classified them according to the Dijkstra (2013).

### 2.3: Species richness and habitat association

We calculated the species richness of each study site by the number of odonates in each sites. We classified the habitat of the study site into six different classes such as Semi Urban (SU), Marsh Lands (ML), Riverside (RS), Lakeside (LS), Forest Patches (FP) and Mangrove Forest (MF). Finally, we calculated species richness in each habitat class and determine the relative species abundance in those habitat types.

### 2.4: Correlation between species richness and climatic variables

We collected monthly average temperature, maximum rainfall and relative humidity data of Khulna region from Bangladesh Bureau of Statistics. We calculated the pearson correlation between species richness and those climatic variation in GraphPad Prison 6.0.

## 3. Results

### 3.1 Species richness in study area

In total 50 species of 30 genera belonging to six families were recorded from the study area. Among them, 31 species (62%) belonging to 22 genera were recorded from Anisoptera sub-order while 19 species (38%) comprises of eight genera were reported from Zygoptera suborder (Table 02). Libellulidae was the most dominant family with 56% (28 species) of the total species count. Coenagrionidae showed next highest dominance with 34% (17 species) species count, followed by Platycnemididae (4%), Protoneuridae (2%), Gomphidae (2%), Aeshnidae (2%) and Corduliidae (2%). Libellulidae was the best represented Anisopteran family with 28 species whereas Coenagrionidae was the most abundant Zygopteran family with 17 species. Agriocnemis is recorded as most dominated genera with 5 species, Orthetrum contribute 4 species, Ceriagrion, Brachydiplax, Ischnura contributed 3 species each whereas Diplacodes, Neurothemis, Rhodothemis, Trithemis, Mortonagrion and Pseudagrion contributed 2 species individually and rest of the genera contributed only a single species.

**Table 02.**
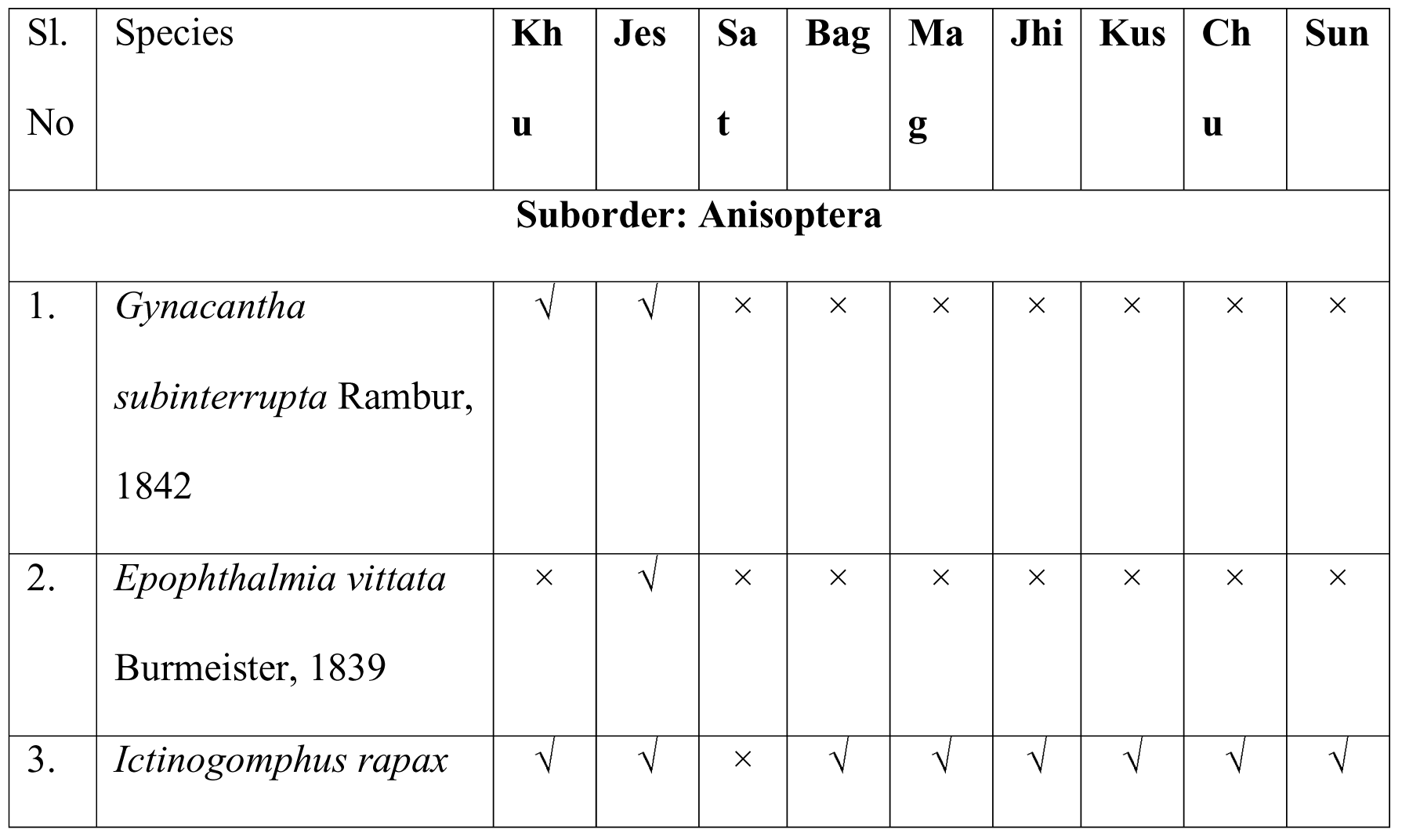
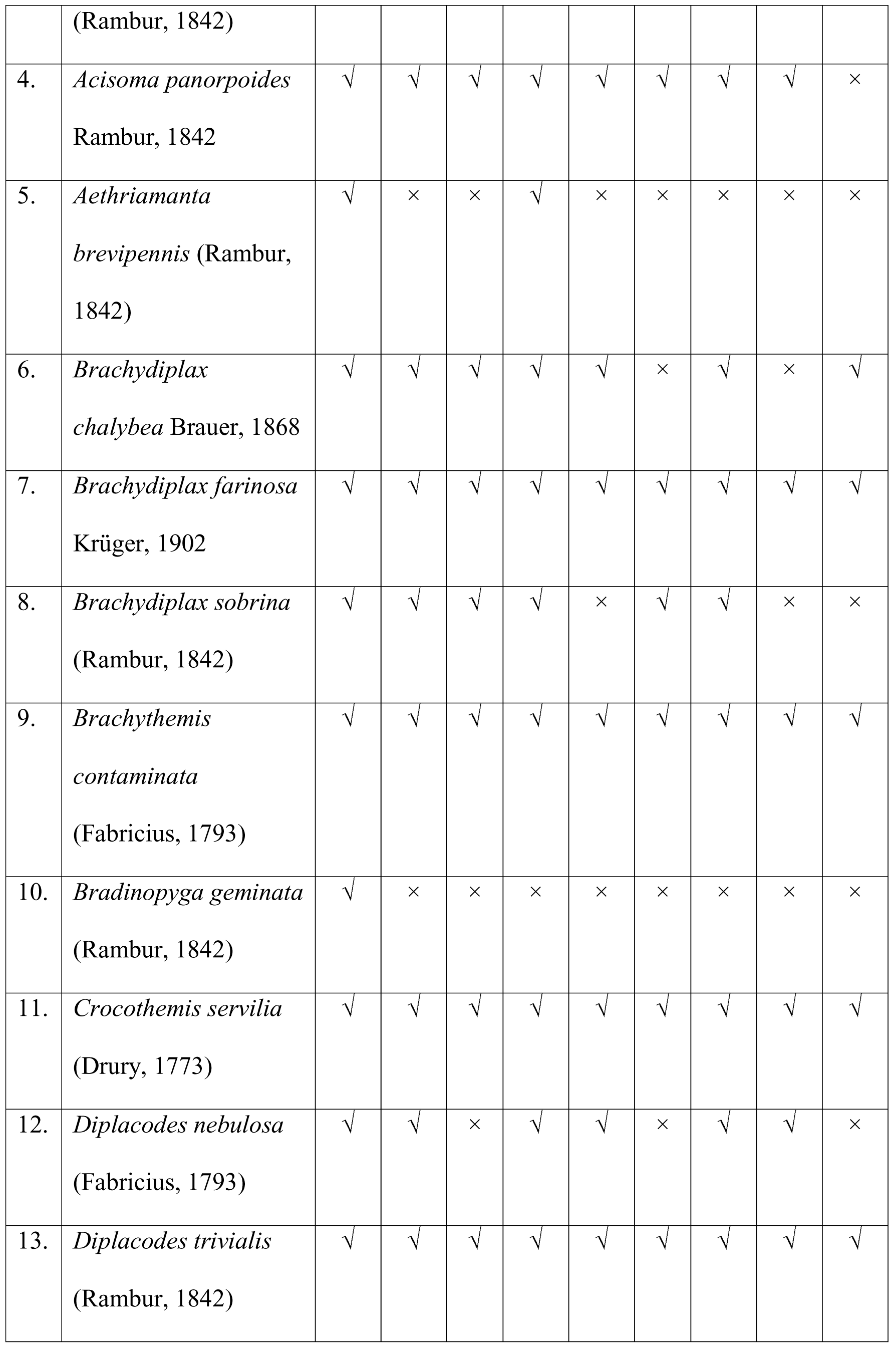
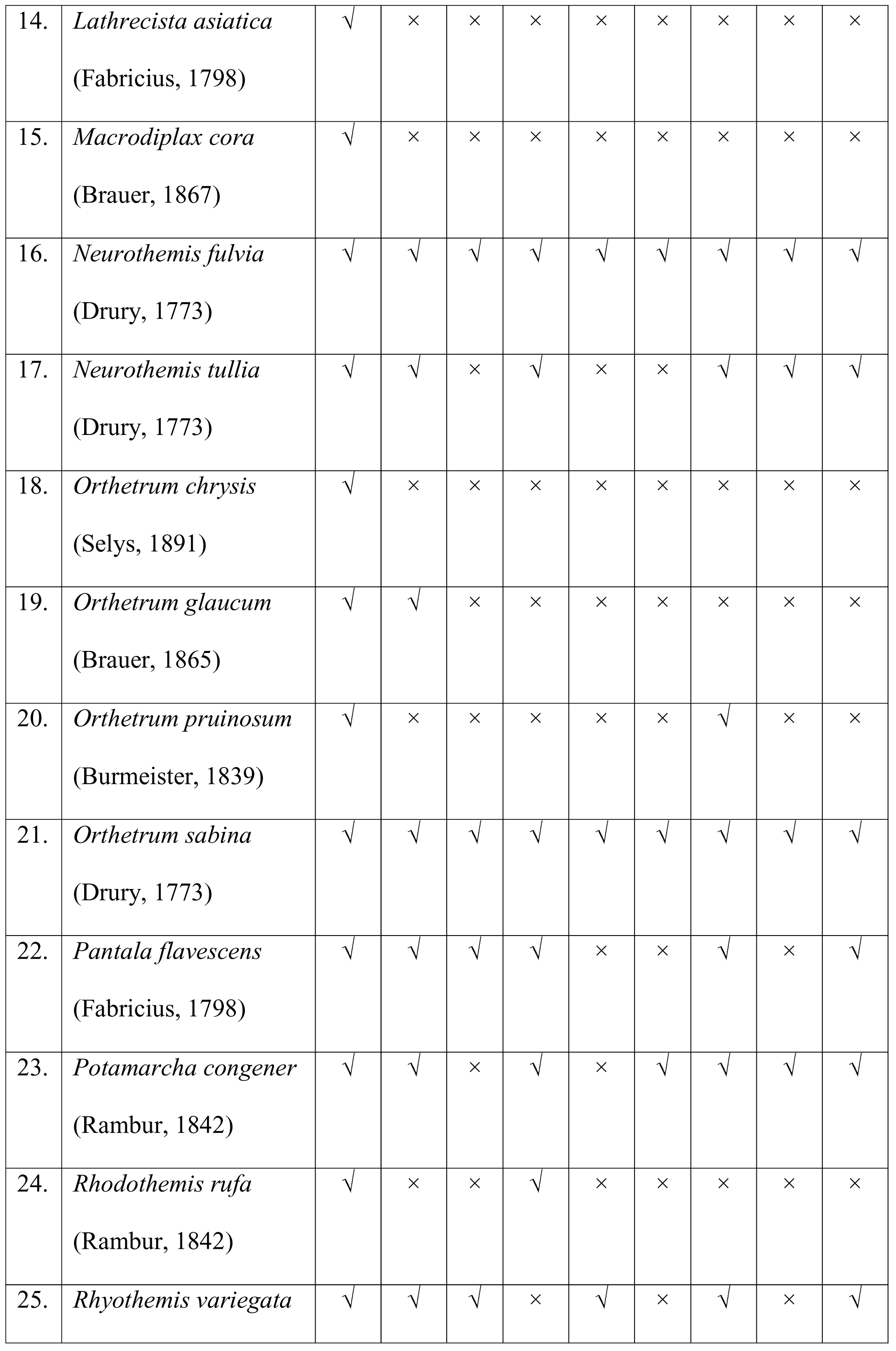
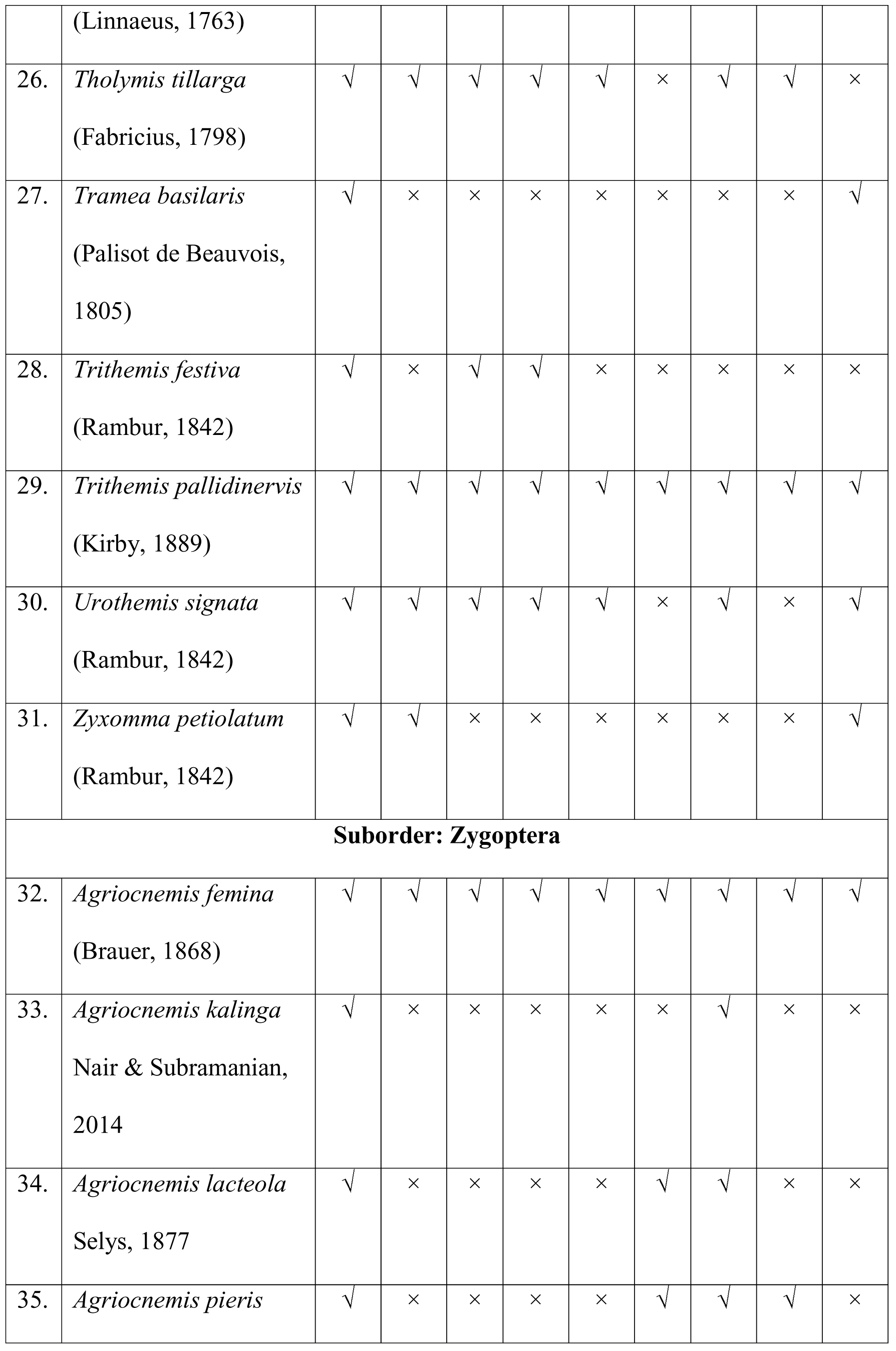
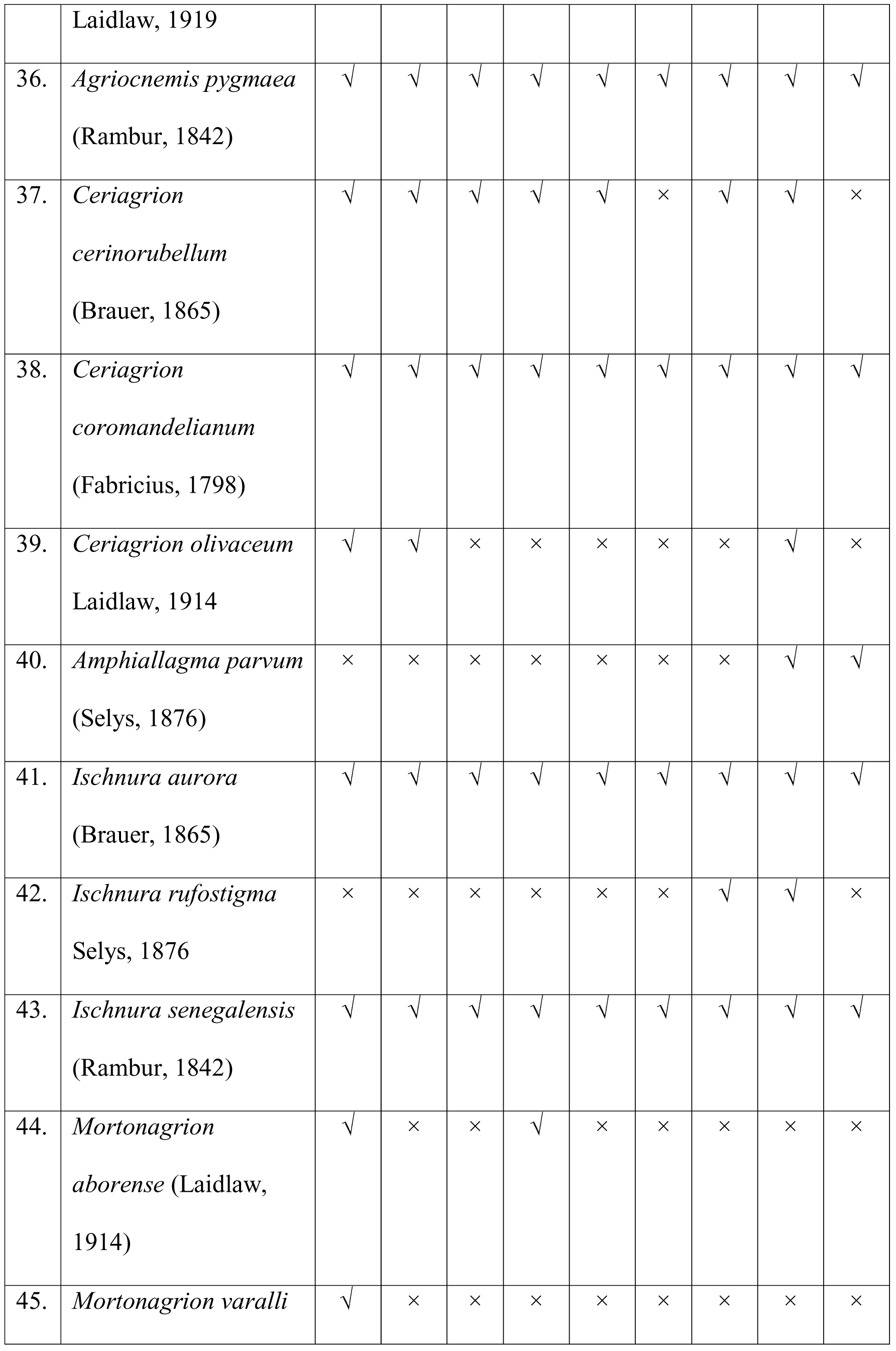
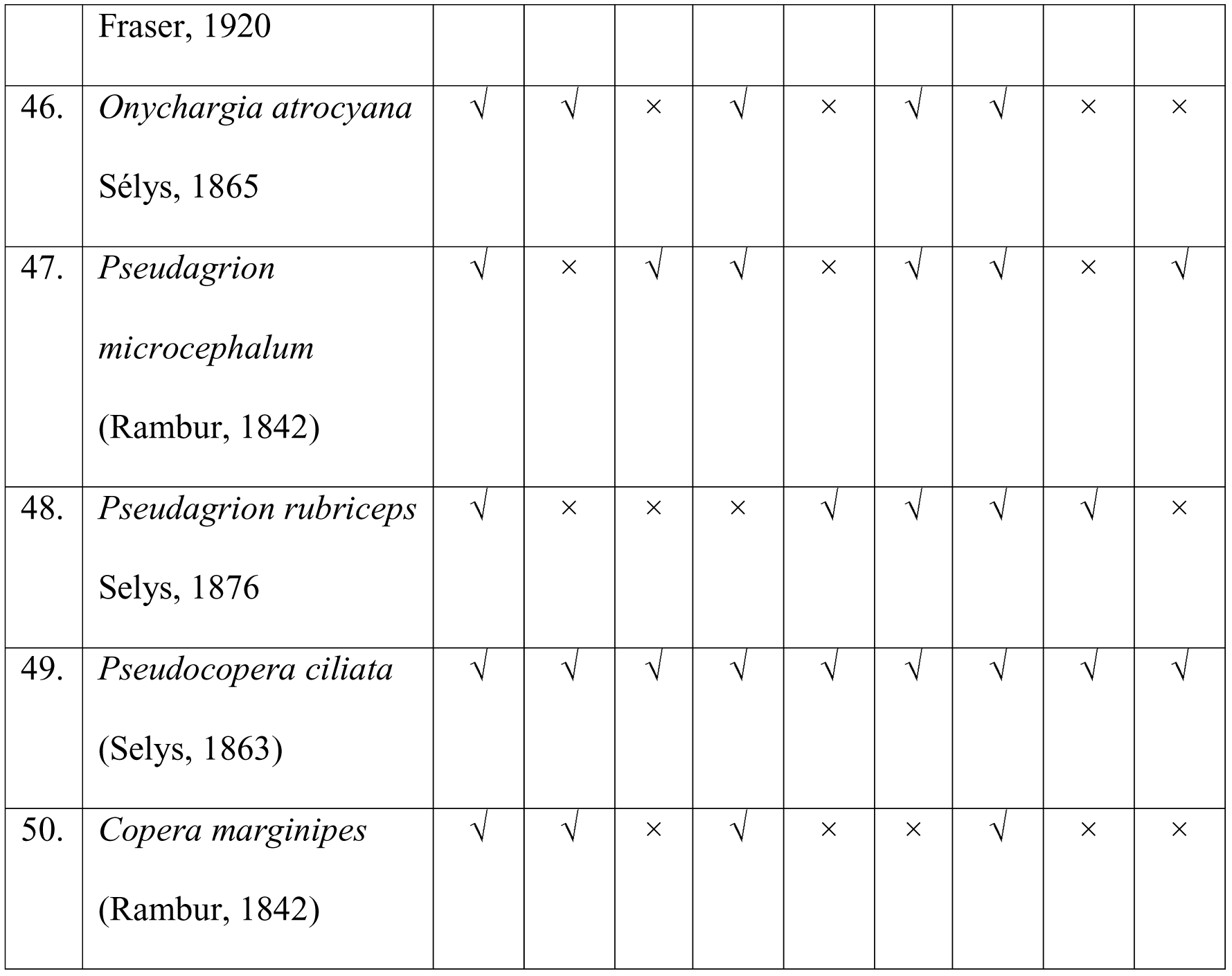
A list of odonates in current study. Khu=Khulna, Jes=Jessore, Sat=Satkhira, Bag=Bagherhat, Mag=Magura, Jhi=Jhinaidha, Kus=Kushita, Chu=Chuadanga, Sun=Sundarbans, √=presence and ×=absence. Species newly added to the Bangladeshi Odonata fauna have been marked with asterisks.

Maximum 47 species were recorded from Khulna district followed by Kushita (36), Jessore (32), Bagherhat (32), Chuadanga (27), Satkhira (24) Magura (22) and Jhinaidha (22). A total of 25 species were recorded from the Sundarbans Mangrove area (Fig 03). All 45 selected survey sites have different species composition however 13 species were commonly found in all zones/districts of the study area. In this study, *Agriocnemis femina, Agriocnemis pygmaea, Brachydiplax farinose, Brachythemis contaminate, Ceriagrion coromandelianum, Pseudocopera ciliata, Crocothemis servilia, Diplacodes trivialis, Ischnura aurora, Ischnura senegalensis, Neurothemis fulvia, Orthetrum sabina,* and *Trithemis pallidinervis* are found in all districts or zones of Khulna division. *Agriocnemis pygmaea, Diplacodes trivialis* and *Ischnura senegalensis* are the most dominated species found in this study. This three species were recorded from all 45 surveyed sites. *Bradinopyga geminate, Amphiallagma parvum, Lathrecista asiatica* and *Orthetrum chrysis* are rare in the study region and recorded from a single site only (Table 02). *Mortonagrion varalli* Fraser, 1920 was first time recorded from Bangladesh from a single female collected from the Khulna University campus.

### 3.2 Habitat association of the odonata

Among the six classified habitat types, marshland has highest species richness with 38 species followed by 34 species in the lake side (Fig.2). On the other hand forest patches, semi urban, riverside habitat type shows similar species richness with 31, 29 and 27 species respectively (Fig. 2). We have found the list number of species at mangroves area, which consists with 25 species (Fig. 2).

**Figure 2:**
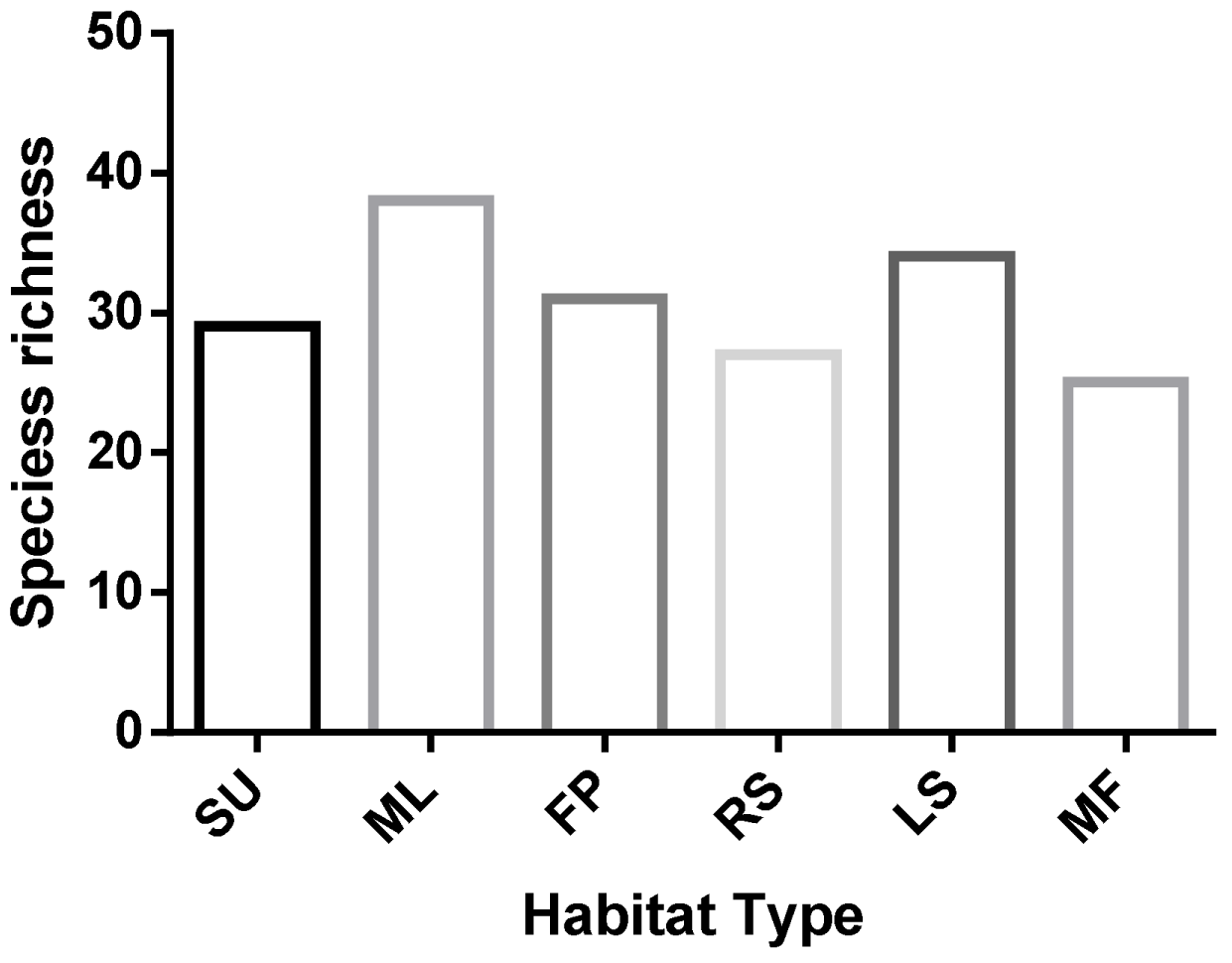
Species richness in different habitat type of the south-western region of Bangladesh.

### 3.3 Correlation of the species richness and abiotic factor

Species richness is positively correlated with the monthly average temperature (p<0.001), relative humidity (p<0.0001) and maximum rainfall (p<0.0001) (fig. 3A, 3B, 3C). Monthly distribution of species richness shows a bell shape curve (fig. 4). Species richness is lowest from December to April (Fig. 4). After April, species richness gradually increases and reached its peak in August. After August species richness again starts to decreases and riches is lowest in between December-February.

**Figure 3:**
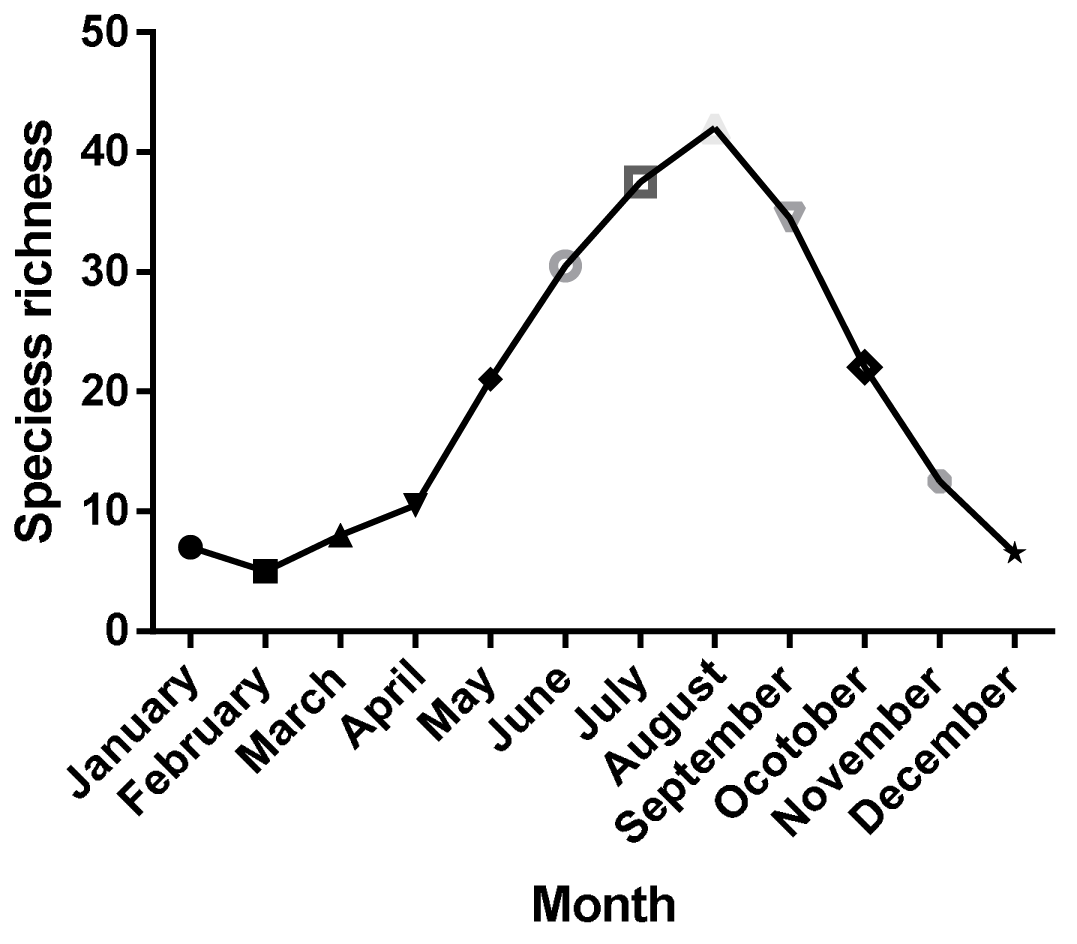
Average monthly species richness in different months.

**Figure 3:**
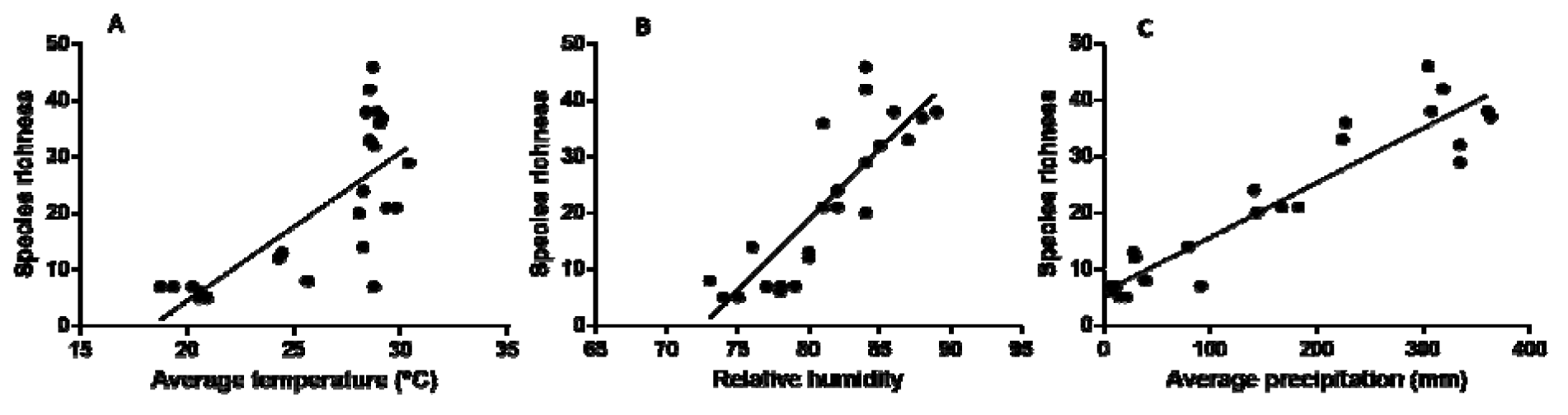
Correlation between A) species richness and average temperature B) Species richness and relative humidity and 3) temperature vs average precipitation

## 4 Discussion

We have updated the checklist of the south-west region of Bangladesh which consists 50 species now and added a news species to the current checklist of the dragonflies and damselflies of Bangladesh. We have also shown that species richness varies in different habitat and highest in marshlands. Our study also shows that, species richness is also influenced by seasonal variation such as temperature, rainfall and humidity.

As a semi-aquatic insect, odonates depend heavily on the waterbodies. Several previous studies showed odonata diversity is high close to the waterbodies (Hornung and Rice 2003; Oppel 2005; De pavia Silva et al. 2010). In our study we have found the similar pattern by discovering most of the species at marsh land and lake side habitats. On the other hand, species richness were lowest in the Mangrove forest area. The water salinity of the Bangladeshi mangroves varies 5ppt to 25ppt (Joshi and Ghose 2003). Water quality is also very important factor for odonata richness (Carchini et al. 2005) and salinity water have negative influences on the odonatan diversity (ref). So the lower, odonata diversity in the mangrove area can be caused by the water salinity. However, larval studies in this region with varies water salinity is needed to understand the impact of salinity on odonata diversity.

Our results show that, species richness in the study area is positively correlated with the rainfall, humidity and temperature. Dragonflies depends on the water for their larval development. After mating, the females lay eggs on the water, or submerged water. Hence, species richness is highest in the monsoon and after monsoon when rainfall is maximum and water reservoirs are full with water. Dragonfly larva require an ambient temperature to emerge in adults (Corbet, 1999). Moreover, like other ectotherm, odonates depend on the external temperature to get ambient body temperature for flight. In congruent to that, our study also shows high species richness in summer and monsoon when temperature is high.

At the current study we have recorded 50 species from the south western region of Bangladesh which is 47% of the total country documented species in Bangladesh. We have discovered one species new for Bangladesh. The newly added damselfly to the Bangladeshi fauna, *Mortonagrion varalli,* was recorded during our regular sampling period from Khulna University campus. On the other hand, another rare dragonfly *Epophthalmia vittata* was recorded during opportunistic survey. In our study, the regular survey was done to survey sites of Khulna and Jessore covering the all seasons but other sites were surveyed in opportunistic survey. Similar to previous study the current study also suggests opportunistic survey can be important for observing odonata species (Koparde et al. 2014, Khan, 2015b,).

With the change of habitat and microclimate, the species content varies. Different types of species found in different habitats. Some of the species found distinctly habitat specialist but some are behaving as generalist. *Agriocnemis femina*, *Agriocnemis kalinga*, *Agriocnemis pygmaea*, *Diplacodes nebulosa* and *Diplacodes trivialis* are found more or less restricted to grassland habitats. *Mortonagrion varalli*, *Ischnura senegalensis*, *Pseudagrion microcephalum* and *Pseudagrion rubriceps* are very close to water bodies which indicates high influence of water to them. Beside them, *Epophthalmia vittata*, *Lathrecista asiatica* and *Gynacantha subinterrupta* are found only at the higher canopy of certain places.

Among six recorded families, species recorded from Aeshnidae and Corduliidae are lower in comparison to the other families. Only a single species of those two families were sighted from the study area. This indicate the rarity of this species in this region of Bangladesh. Platycnemididae has contributed with two species. From this two, *Pseudocopera ciliate* were frequently sighted in different study sites. Similar to previous studies (Khan 2015b; Chowdhury 2011), Libellulidae and Coenagrionidae were recorded as the most dominated Anisopteran and Zygopteran families respectively. Except *Lathrecista asiatica* and *Bradinopyga geminate,* all other Anisopteran species were frequently observed. Similarly, Coenagrionidae species were frequent recorded except *Aciagrion* and *Mortonagrion* species.

In conclusion, we have recorded 50 species of dragonflies and damselflies during current survey and updated the checklist of the south-western region of Bangladesh. We have shown species assemblage is higher in the marshland and lakeside however lowest in the mangrove regions. We also shown species richness is correlated with the rainfall, temperature and relative humidity and in the study regions species richness is highest during and after monsoon. Future studies is required to understand the biology and threat of the species in the study area.

